# Fair molecular feature selection unveils universally tumor lineage-informative methylation sites in colorectal cancer

**DOI:** 10.1101/2024.02.22.580595

**Authors:** Xuan Cindy Li, Yuelin Liu, Alejandro A. Schäffer, Stephen M. Mount, S. Cenk Sahinalp

## Abstract

In the era of precision medicine, performing comparative analysis over diverse patient populations is a fundamental step towards tailoring healthcare interventions. However, the critical aspect of equitably selecting molecular features across multiple patients is often overlooked. To address this challenge, we introduce `FALAFL` (FAir muLti-sAmple Feature seLection), an algorithmic approach based on combinatorial optimization. `FALAFL` is designed to bridge the gap between molecular feature selection and algorithmic fairness, ensuring a fair selection of molecular features from all patient samples in a cohort.

We have applied `FALAFL` to the problem of selecting lineage-informative CpG sites within a cohort of colorectal cancer patients subjected to low read coverage single-cell methylation sequencing. Our results demonstrate that `FALAFL` can rapidly and robustly determine the optimal set of CpG sites, which are each well covered by cells across the vast majority of the patients, while ensuring that in each patient a high proportion of these sites have good read coverage. An analysis of the `FALAFL`-selected sites reveals that their tumor lineage-informativeness exhibits a strong correlation across a spectrum of diverse patient profiles. Furthermore, these universally lineage-informative sites are highly enriched in the inter CpG island regions.

`FALAFL` integrates equity considerations into the molecular feature selection from single-cell sequencing data obtained from a patient cohort. We hope that it will help propel equitable healthcare data science practices and contribute to the advancement of our understanding of complex diseases.

## 1 Introduction

The pursuit of precision medicine in modern healthcare has ushered in a promising era of tailored interventions and therapies, where the molecular characteristics of individual patients guide treatment strategies. Central to this pursuit is the intricate task of molecular feature selection, a pivotal process for discerning distinctions and commonalities among patient cohorts. However, a critical aspect that is often overlooked is the fair selection of molecular features across patient samples, especially when patients are subjected to single-cell sequencing with varying read depth. Achieving balance between patients in molecular feature selection is pivotal to prevent bias and ensure interpretability of comparative analysis across patients in a cohort.

Prior research has recognized the importance of fair feature selection, not only in healthcare but across various domains. In natural language processing, [Rios and Bispo, 2018] show that it is crucial to have fair feature selection in text classification tasks to prevent individual terms from dominating the classification process and thereby preserve the integrity and fairness of the results. [Khojasteh et al., 2023] has applied a data balancing technique derived from One-class Support Vector Machine onto drug-target interaction data to enhance prediction of drug-target interactions. These diverse applications point to the sweeping need for fair feature selection methodologies, which aim to maintain balance in the selection process and avoid the undue influences of any single feature or group of features. The balance, in turn, contributes to the robustness and fairness of the analysis and enhances the quality of the gained insights.

The problem of fair molecular feature selection across patient samples is related (but somewhat orthogonal) to the well-known “hitting set problem [Karp, 1972].” The hitting set problem seeks to find the smallest possible set of elements *H* (referred to as the “hitting set”) from a given collection *C* of sets *C*_1_, *C*_2_, … such that at least one element *c*_*i,j*_ from each set *C*_*i*_ is included in *H*. In other words, this problem asks to identify a minimal set *H* that “hits” or intersects with each of the sets *C*_*i*_ in the collection *C*.

The molecular feature selection problem, on the other hand, has a similar setting but a very different objective. In this problem, each set *C*_*i*_ for a given patient *i* is composed of elements *c*_*i,j*_, each of which corresponding to a distinct molecular feature, such as a genetic marker, expression of a gene, or an epigenetic characteristic. To select a rich and balanced set of molecular features, one needs to come up with the *largest* subset of molecular features such that not only each feature is present in at least a user defined number of patients, but also each patient contributes equitably to the set of selected features. In real-world applications, the molecular feature selection problem is further complicated by varying read depth of the samples, especially in cases involving single-cell sequencing for each patient. For instance, in a colorectal cancer (CRC) single-cell dataset published by [Bian et al., 2018], there is significant variability in the number of cells collected from each patient and the coverage of CpG sites in each cell (see Figure S1). This observed inconsistency within the same cohort complicates the attempt to select CpG sites that adequately represent the entire cohort, thus muddling the downstream efforts of understanding common epimutation dynamics across patients and identifying driver methylation events. These real-world challenges call for a systematic approach to fairly select molecular features that effectively encapsulate the entire patient cohort.

To overcome the said obstacles, in this paper, we introduce `FALAFL` (FAir muLti-sAmple Feature seLection), a novel fair molecular feature selection scheme grounded in integer linear programming (ILP). Designed to introduce algorithmic fairness in comparative patient profile analysis, the primary goal of `FALAFL` is to provide a fast and reliable solution for the fair selection of molecular features across different patient samples to represent the entire patient cohort. When finding the optimal solution for feature selection, `FALAFL` not only achieves balance across all patients, but also ensures that each patient receives sufficient representation.

We have applied `FALAFL` to the problem of fair selection of CpG sites within the aforementioned CRC cohort [Bian et al., 2018], in which the patients are subjected to low coverage single-cell methylation sequencing. We demonstrate that `FALAFL` rapidly and robustly identifies a maximal set of CpG sites that are well-represented across a majority of patients, while ensuring that each patient contributes a proportionate number of sites to the final selection, enabling equitable comparative analysis of the selected sites. We then performed such a comparative downstream analysis of the `FALAFL`-selected sites, which revealed that lineage-informativeness among the CpG sites, i.e., the property that the metylation status of a CpG site is stably inherited in tumor progression, is universal. These findings unveil the universal non-stochastic nature of methylation changes in the course of colorectal cancer progression.

## 2 Method

The input to `FALAFL` is genomic, epigenomic or transcriptomic sequencing data collected from a diverse set of tumor samples. For the remainder of the paper, we will focus on single-cell methylation sequencing data, however `FALAFL` is applicable to other types of sequencing data as well. In single-cell methylation sequencing, the average read depth across any given cell’s genome is not only very low (0.1 × or lower), but also highly variable across the CpG sites, cells and samples. Nevertheless, because methylation statuses of many CpG sites are altered during tumor evolution, there has been attempts to reconstruct a tumor’s progression history (i.e. tumor phylogeny) by the use of single-cell methylation sequencing data [Bian et al., 2018]. Unfortunately, not all CpG sites are in a tumor are “lineage-informative”, i.e. could be employed in the reconstruction of a tumor phylogeny, since their methylation statuses are either static or are too dynamic to be associated with distinct lineages. It is, however, possible to identify these lineage-informative sites in a tumor sample, while simultaneously inferring the tumor phylogeny, by an iterative approach named `Sgootr` [Liu et al., 2023]. `Sgootr` has been applied to the single-cell methylation sequencing dataset from several colorectal cancer patients [Bian et al., 2018] to demonstrate that the number of distinct metastatic seeding events in these tumors are fewer than those reported in the original study. However, because `Sgootr` only uses CpG sites that are well covered across the cells of a given tumor sample, the set of CpG sites `Sgootr` identified as lineage-informative in one sample can be quite different from those identified as lineage-informative in another sample. This makes it difficult to assess whether lineage-informativeness is a universal property of CpG sites across all or a subset/subtype of colorectal cancers, or is tumor specific. Such an assessment is important since universality in lineage-informativeness could imply a functional association between lineage-informative CpG sites and colorectal cancer evolution. It could also provide means of evaluating the clonality and advancement of a tumor.

As we describe below, `FALAFL` takes as the input *the read depth information*^1^ for each CpG site, in each cell, from each tumor sample, and outputs a set of CpG sites, each with sufficient read depth across the cells of nearly all (i.e. with the exception of a small number of) patients. Importantly, the `FALAFL` identified CpG sites are also balanced across tumor samples, in the sense that, the proportion of these CpG sites with sufficient read depth across a large proportion of cells in each tumor sample will be higher than a user defined threshold. On these balanced selection of CpG sites, one can evaluate lineage-informativeness in each tumor and assess whether this property is truly universal in colorectal cancer or tumor specific.

### 2.1 FALAFL: fair selection of molecular features for multi-patient comparative analysis

The input for `FALAFL` is a multi-patient integrated CpG site read coverage data represented as a patient-by-site matrix *S*_*n*×*m*_, where *n* is the number of patients, *m* the number of CpG sites, and *s*_*i,j*_ the fraction of cells in patient (i.e. tumor sample) *i* in which CpG site *j* has “sufficient” read depth (e.g. two reads or more) as defined by the user.

In a first preprocessing step, we ensure that for each patient *i*, we have *s*_*i,j*_ *> δ* (for some user defined *δ*) for each site *j*; if not, site *j* is not considered any further for that patient. (This preprocessing step is to reduce the size of input matrix.)

In a second preprocessing step, we binarize *S* to obtain 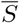, where 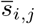 indicates whether site *j* has sufficient read depth in at least a fraction of *p* cells in patient *i*. (This second step is to conceptually simplify the problem.)

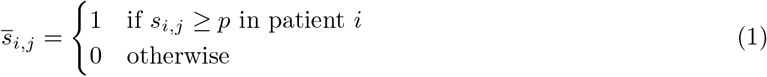

In a third preprocessing step, we eliminate all sites *j* where the total number of patients *i* with *s*_*i,j*_ = 1 is ≤ *k* (for some user defined threshold *k*). (This step also aims to reduce the size of the problem.) We achieve this by removing all sites *j* for which 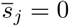:

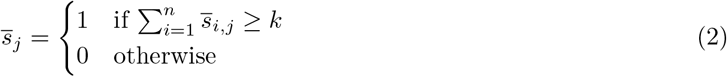

Now we can define our combinatorial optimization formulation on the preprocessed binary input matrix 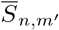 (where *m*^′^ denotes the number of sites not eliminated in the preprocessing steps), given a user defined threshold *q*, which denotes for each patient *i*, the minimum proportion of sites *s*_*j*_ chosen by the formulation for which 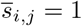. Let **r** be a binary vector where *r*_*j*_ = 1 indicates that site *j* is chosen by the formulation. Then our formulation chooses sites with the objective to maximize:

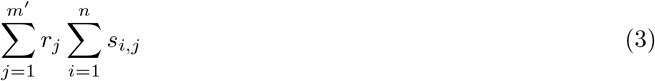

subject to the constraint that:

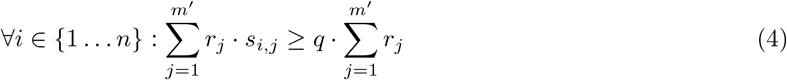

The above formulation of `FALAFL` is implemented using the Gurobi ILP solver [Gurobi Optimization, LLC, 2022]. The code availability of `FALAFL` can be found in Section S1.

### 2.2 Evaluating lineage-informativeness of FALAFL-selected CpG sites from colorectal cancer patients by using tumor phylogenies

A single-cell methylation sequencing (more specifically, single-cell bisulfite sequencing, or sc-BSseq) dataset has been made available by [Bian et al., 2018] for nine metastatic colorectal cancer patients. A description of the number of cells, number of input CpG sites, as well as lesion information for each patient can be found in Table S1. The distribution of the number of CpG sites sequenced per cell in each patient can be found in Figure S1. As will be described in detail in Section 3.1, we apply `FALAFL` to the single-cell methylation sequencing data from these 9 patients to identify ∼ 196*K* CpG sites, which are well covered across the cells of a large fraction of patients, while ensuring that a large fraction of these sites are well covered in each of the patients. (Separately, we also apply`FALAFL` to identify ∼ 1.35*M* CpG sites from 4 patients with a primary tumor in the left colon; these patients have better cell coverage and read depth than the others.) On these CpG sites, we evaluate the lineage-informativeness in each patient as described below.

For each patient, `Sgootr` [Liu et al., 2023] is applied to the input read count matrices *N* (the matrix with methylated read counts) and *M* (the matrix with unmethylated read counts), in each of which the entry at row *i* and column *j* represents the read count of the CpG site *i* in the cell *j*, for obtaining a tumor phylogeny. `Sgootr` computes the tumor phylogeny iteratively: in each iteration, it prunes a fraction of lineage-uninformative CpG sites and builds the tumor phylogeny with the unpruned CpG sites. We set the the pruning fraction *κ* to .1 (the default setting). This means that in each iteration, the “least lineage-informative” 10% of the CpG sites (which have been used to construct the tumor phylogeny in the previous iteration) are pruned out. Note that the lineage-informativeness of a CpG site is determined by its methylation persistence score defined in Equation 6 in [Liu et al., 2023], which is based on the methylation status of the site across the subtrees of the tumor phylogeny obtained in the previous iteration. More specifically, the methylation persistence of a CpG site and a given node of the tumor phylogeny is the Jensen–Shannon (JS) distance [Wong and You, 1985, Lin, 1991] between the site’s methylation rate distribution in the subtree rooted at that node, and that of the remainder of the tumor phylogeny. A high JS distance indicates that the CpG site’s methylation status across the leaves of the subtree is significantly different from that of the leaves in the remainder of the tumor phylogeny, i.e. its methylation status change at the aforementioned node of the subtree and this change is stably inherited by the descendants of this root node. The lineage-informativeness of the CpG site for a given tumor phylogeny is then defined to be the maximum methylation persistence of that site across all internal nodes of the tumor phylogeny.

It is important to note that the iterative process of `Sgootr` terminates after a user defined number of rounds (whose default setting is 40). Among the tumor phylogenies obtained in these iterations, `Sgootr` outputs the tree which is “most stable”, i.e., whose Robinson-Foulds (RF) distance [Robinson and Foulds, 1981, Sul and Williams, 2008] with the tree obtains in the next iteration is the smallest. The lineage-informativeness of each CpG site is calculated on this tumor phylogeny outputted by `Sgootr`.

### 2.3 Evaluating universality of lineage-informativeness using deviation from perfection correlation

The lineage-informativeness of a site *j* across all *n* patients can be represented as a vector **v**_**j**_ of size *n*:

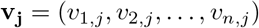

For the sake of brevity, we simplify the notation of vector **v**_**j**_ as **v** = (*v*_1_, *v*_2_, …, *v*_*n*_). Here, 0 ≤ *v*_*i*_ ≤ 1, since lineage-informativeness is measured based on the JS distance as described earlier, which ranges between 0 and 1.

The universality of lineage-informativeness of CpG site *j* across all *n* patients can thus be evaluated as the *L*_2_ distance between the point *V* ∈ ℝ^*n*^ with coordinates (*v*_1_, *v*_2_, …, *v*_*n*_) (which corresponds to the vector **v**), and the line *ℓ* of perfect equality, between the points *ℓ*_**0**_ = (0, 0, …, 0) and *ℓ*_**1**_ = (1, 1, …, 1), where *ℓ*_**0**_, *ℓ*_**1**_ ∈ ℝ^*n*^. Note that if the CpG site is truly universal, i.e. has identical methylation persistence across all patients, then **v** would be on the line of perfect equality.

Let **n** be the unit vector parallel to *ℓ*, i.e., 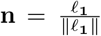, then the *L*_2_ distance between *V* and *ℓ* can be computed as follows:

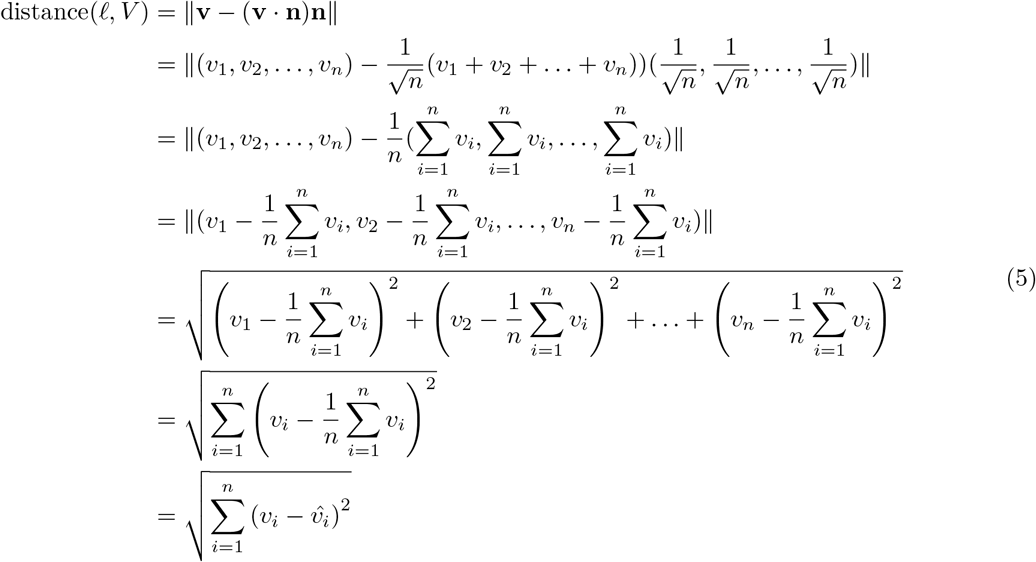

Thus, distance(*ℓ, V*) is equivalent to the mean square difference of vector **v**. We normalize the resulting distance using the maximum distance possible in the *n*-dimensional Euclidean space confined by 0 ≤ *v*_*i*_ ≤ 1 in the *i*th dimension for all *n* dimensions; this is because we would like to use a measure that is independent from the number of patients - which will become important when comparing the universality of lineage-informativeness across all nine patients with that among those patients with each specific tumor subtype.

The mean square difference is maximized when the coordinates of *v*_*i*_ are as disperse as possible, i.e., half of *v*_*i*_ needs to be 0 and the other half 1. If *n* is an even number, the maximum mean square difference of vector **v** is 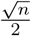. If *n* is an odd number, the maximum mean square difference of vector is then 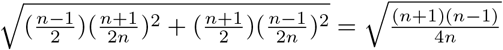. Therefore, the maximum distance(*ℓ, V*) is the following:

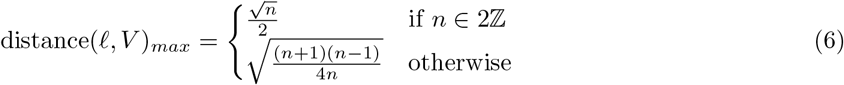

Thus, the normalized distance, which we call *deviation from perfect correlation*, is the following:

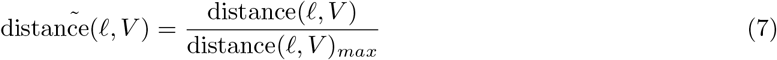

### 2.4 Defining universally lineage-informative and uninformative CpG sites

As aforementioned, the lineage-informativeness of a CpG site characterizes how stably inherited its methylation change is in a given patient, and the universality of lineage-informativeness measures how conserved its methylation change dynamics is across different patients. Here, we formally define universally lineage-informative and uninformative CpG sites. In essence, the universality of a site’s behavior across patients is determined by its deviation from perfect correlation, as described in Section 2.3, while lineage-informativeness is determined by the mean JS distance of the site across patients. Universally lineage-informative sites have a large mean JS distance across all *n* patients, but exhibit only a small deviation from perfect correlation. Conversely, universally uninformative sites display both a small mean JS distance across all patients and a small deviation from perfect correlation.

Specifically, we deem a CpG site universally lineage-informative, if its mean JS distance exceeds the mean value of all sites’ mean JS distances across patients, and its deviation from perfect correlation is smaller than the mean perfect correlation of sites. On the contrary, if a site’s mean JS distance is smaller than the mean value of all sites’ mean JS distances across patients, and its deviation from perfect correlation is smaller than the mean perfect correlation of all sites, then it is classified as universally uninformative.

### 2.5 Greedy pairwise selection and random selection of CpG sites

We use greedy pairwise selection and random selection of CpG sites to benchmark against `FALAFL`-selected CpG sites specifically on the subcohort of 4 patients with the primary tumor in the left colon, primarily because these patients have higher numbers of cells and better read depth coverage. Additionally, we would like to perform the benchmarking analysis within a homogenous patient cohort to avoid potential perturbations due to high levels of patient diversity. The greedy pairwise selection seeks to select CpG sites to represent a given patient pair. For each patient pair, sites covered by at least one read in 50% or more of the cells of both patients are selected. In random selection, sites of equal number as `FALAFL` output for the 4 patient subcohort (i.e., 1,346,130 sites) is randomly chosen from the `FALAFL` input matrix, which is sized at 4 × 27, 227, 230. The random selection is repeated 5 times.

## 3 Results

We apply `FALAFL` to the scBS-seq dataset generated by Bian *et al*. on 9 metastatic colorectal cancer patients^2^. In this cohort, patient CRC01, CRC10, CRC11, and CRC13 have primary tumor sites in the left colon, CRC02, CRC04, and CRC15 in the right colon, and CRC12 and CRC14 in the rectum. Here, we present results generated from applying `FALAFL` to the entire CRC cohort as well as the subcohort of 4 patients whose primary tumor is in the left colon (i.e. left colon CRC subcohort). We also include the results obtained from benchmarking `FALAFL` against the greedy pairwise selection and random selection approaches described in Section 2.5 for the left colon CRC subcohort.

### 3.1 FALAFL performs fast and robust molecular feature selection

We prepare the patient data from the left colon CRC subcohort as well as the entire patient cohort for preprocessing into the input matrix tailored for `FALAFL`. For both cohorts, we choose a value of *δ* = 0.1, indicating that a CpG site needs to be covered by at least one read in at least 10% of the cells in each of the patients.

Specific to the left colon CRC subcohort of 4 patients, we configure the input parameters for preprocessing as *p* = 0.5 and *k* = 2, indicating that a CpG site can be considered by `FALAFL` only if it has at least one read covering it in more than 50% of the cells in two or more patients. This results in the generation of two input matrices of the same dimensions, denoted as *S* and 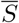, as described in Section 2.1. For the left colon CRC subcohort, the input matrices are sized at 4 × 27, 227, 230. We choose *q* = 0.75 as the input parameter for the ILP constraint as defined in Equation 4, which indicates that in the output matrix, each patient needs to have sufficient coverage in at least 75% of the `FALAFL`-selected sites.

Specific to the entire CRC cohort of 9 patients, we use *p* = 0.5 and *k* = 4 as the input parameters for preprocessing. As the 9 patients have different primary tumor sites and sequencing coverage, we further add a constraint that each CpG site must have coverage in at least 2 left colon CRC patients, 1 right colon CRC patient, and 1 rectum CRC patient. This yields two input matrices *S* and 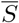 both sized at 9 × 1, 175, 877. The choice for the input parameter for the ILP constraint is *q* = 0.75.

Despite the large dimensionality of the input data, `FALAFL` manages to deliver outputs almost instantly. On a single CPU, `FALAFL` completes feature selection for both the left colon CRC subcohort and the entire CRC patient cohort in less than 60 seconds, underscoring the computational efficiency of our approach. For the left colon CRC subcohort, `FALAFL` yields an output consisting of 1,346,130 selected sites for the 4 patients; for the entire CRC patient cohort, `FALAFL` select 195,809 sites for the 9 patients. For each patient cohort, the application of `FALAFL` on the respective input matrix is repeated three times and the runtime is shown in Figure 2a.

**Figure 1:**
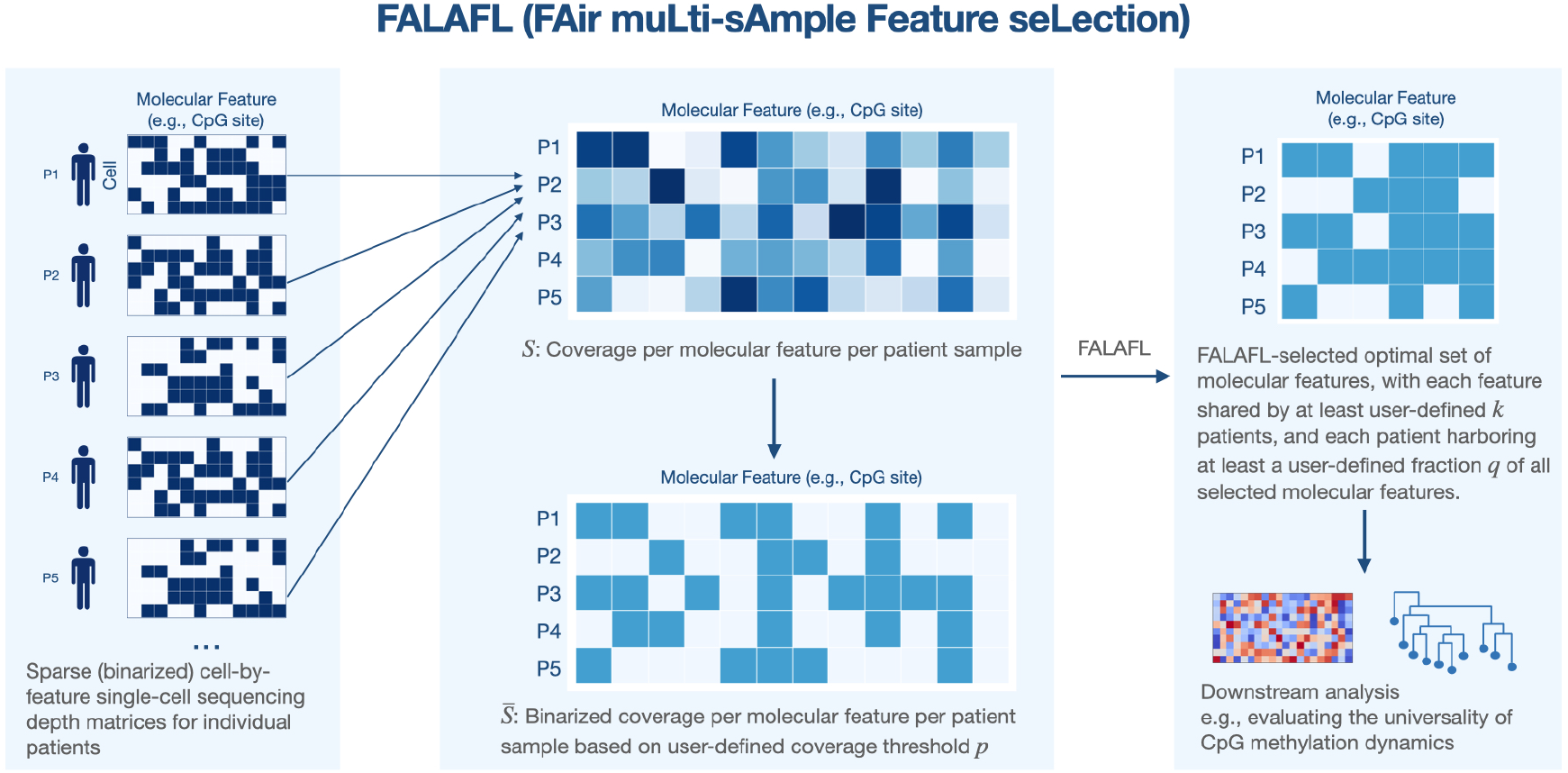
Graphical abstract of FALAFL. `FALAFL` takes the cohort molecular abudance profile and uses integer linear programming to output a set of molecular features that optimally represent the entire cohort. The `FALAFL`-selected features can then be used for downstream analysis, such as understanding the universality of lineage-informativeness of CpG sites in tumor progression.

**Figure 2:**
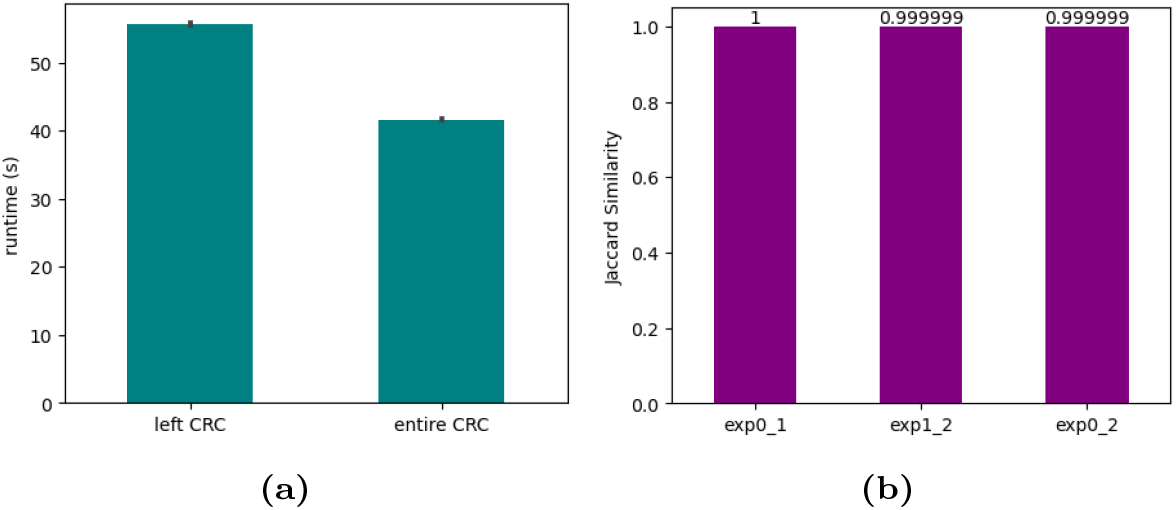
FALAFL runtime and robustness. **(a)** Bar plot showing the `FALAFL` runtime of feature selection for the left colon CRC subcohort and the entire CRC cohort. **(b)** Bar plot showing the Jaccard index of the `FALAFL` output for each pair of the shuffled inputs.

To evaluate the robustness of our output, we perform 3 instances of data shuffling on the input matrix three times for the 4-patient left colon CRC subcohort. More specifically, we randomly shuffle the order of the columns of the input matrix to assess whether `FALAFL` favors certain columns over the others when producing outputs. We compare 3 experiments of data shuffling in pairs: for each pair of the experiments, we evaluate the similarity of the `FALAFL`-outputted sites using the Jaccard index. As indicated in Figure 2b, the results demonstrate a high level of consistency - for all three pairs of the data shuffling experiments, the outputs of `FALAFL` overlap highly, with Jaccard index ranging from 0.9999 to 1.0. The high degree of agreement among the outputs further attests to the reliability and stability of our selection process, signifying the consistency and reproducibility of `FALAFL`’s results.

### 3.2 FALAFL optimizes feature selection for the entire patient cohort

To demonstrate the ability of `FALAFL` to perform fair selection for the entire patient cohort, we benchmark `FALAFL` against the greedy pairwise selection using the data of the left colon CRC subcohort. We perform greedy pairwise selection which maximizes the number of shared sites between each pair of patients in the subcohort, as detailed in Section 2.5. The right panel of Figure 4a provide clear evidence that `FALAFL` balances the number of sites shared across different pairs of patients, whereas the greedy pairwise approach selects drastically different number of sites for each patient pair. Additionally, as shown in the left panel of Figure 4a, `FALAFL` selects a much greater number of sites to represent the left colon CRC subcohort than the greedy pairwise approach does: `FALAFL` outputs ∼ 1.35*M* sites for the left colon CRC subcohort, whereas there are only ∼ 0.47*M* sites shared across all 4 patients selected by the greedy pairwise approach. This result shows that `FALAFL` is able to not only select molecular features to maximally represent the entire patient cohort, but also balance the information shared for each pair of patients without favoring or disregarding any patient pairs.

### 3.3 FALAFL helps discover cohort-wise universal behavior in methylation changes

We use `FALAFL` to select CpG sites representative of the left colon CRC subcohort and the entire CRC cohort, respectively, as described in Section 2.1. We employ the measures described in Section 2.2 and 2.3 to evaluate the universal lineage-informativeness of the `FALAFL`-chosen sites for both cohorts.

As depicted in Figure 3a, the lineage-informativeness level of `FALAFL`-chosen CpG sites for the left colon CRC subcohort are highly correlated among all pairs of patients. As described in Section 2.3, we carry out further examination of the universality of lineage-informativeness levels for all patients in the left colon CRC subcohort. As shown in Figure 3b, the majority of CpG sites display lineage-informativeness levels only slightly deviating from the line of perfect equality, with a median normalized shortest distance of 0.1589. We also evaluate the universality of lineage-informativeness of `FALAFL`-selected CpG sites for the entire CRC cohort of 9 patients. Among the `FALAFL`-selected 195,809 sites, the deviation from perfect correlation is 0.322, as demonstrated in Figure 3c. This result shows that the behaviors of CpG sites are less conservative across a more diverse patient cohort, but nonetheless there are still 11,916 out of the 195,809 `FALAFL`-selected CpG sites with deviations from perfect correlation smaller than 0.1589 (the median deviation for the left colon CRC subcohort). These results manifest the ability of `FALAFL` to discover universally lineage-informative CpG sites in both homogenous patient cohorts and more diverse cohorts.

**Figure 3:**
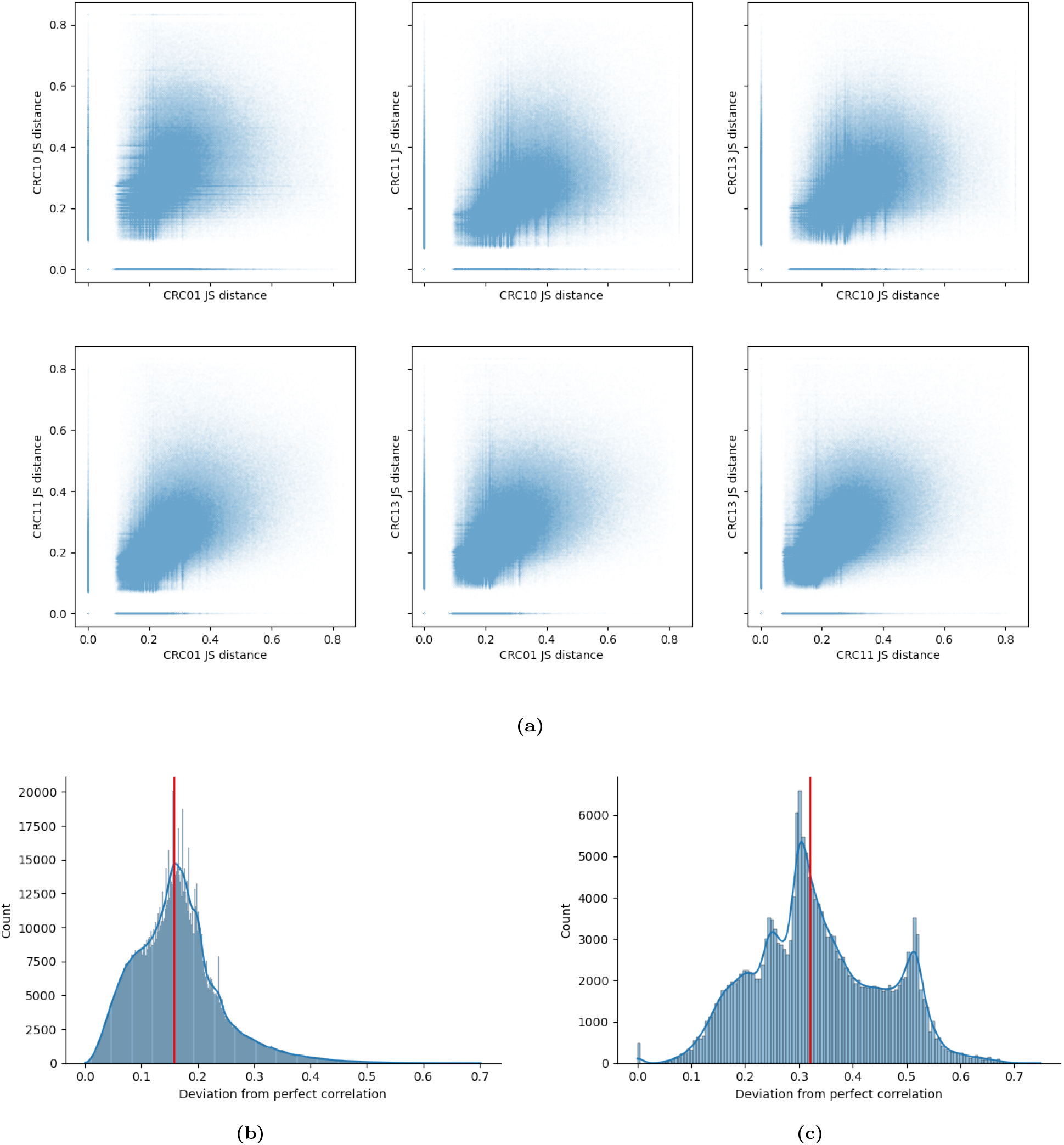
Correlation of lineage-informativeness of FALAFL-chosen CpG sites. **(a)** Pairwise scatter plot showing the lineage-informativeness correlation of `FALAFL`-selected sites for each patient pairs of the left colon CRC subcohort. **(b)** Deviation from perfect correlation for 4 left colon CRC patients. The red vertical line indicates the median deviation from perfect correlation is 0.1589. **(c)** Distance from line of perfect correlation for all 9 patients. The red vertical line indicates the median deviation from perfect correlation is 0.3216

**Figure 4:**
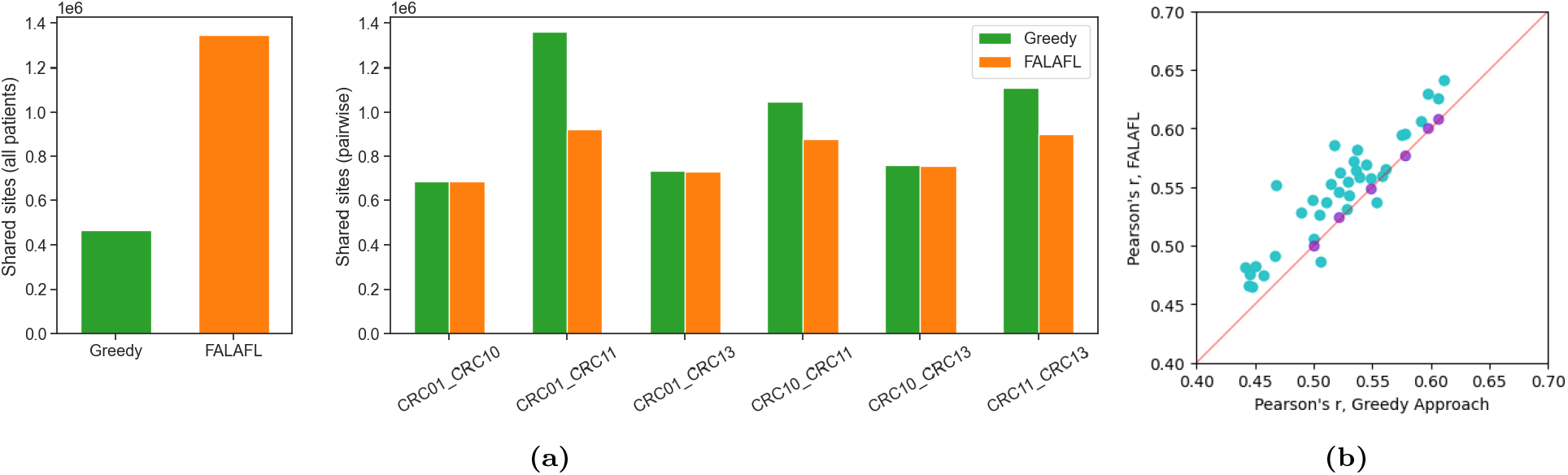
Benchmarking FALAFL against the greedy approach for CpG site selection. **(a)** Comparing the greedy approach and `FALAFL` for fair feature selection. The left panel shows the number of CpG sites shared by all pairs of patients of the left colon CRC subcohort selected by the greedy pairwise approach (effectively selecting sites shared by all patients) vs `FALAFL`-selected sites. The right panel shows the CpG sites present in at least 50% of cells in both patients for each patient pair, selected by the greedy approach vs that by `FALAFL`. **(b)** Comparing the greedy approach against `FALAFL` for correlation of lineage-informativeness for each patient pair. The x-axis is the Pearson’s correlation of lineage-informativeness of CpG sites selected using the greedy approach for each patient pair. The y-axis is the Pearson’s correlation of lineage-informativeness of `FALAFL`-selected CpG sites for each patient pair. The y-coordinates of the purple points is the Pearson’s correlation of lineage-informativeness of CpG sites selected by `FALAFL` in the left colon CRC cohort, measured between each pair of patients. The y-coordiantes of the turquoise points is the Pearson’s correlation of lineage-informativeness of CpG sites selected by `FALAFL` in the entire CRC cohort, again measured for each patient pair. The red line is the diagonal line *y* = *x*.

### 3.4 FALAFL better identifies sites that are universally lineage-informative

We further show that the fair site selection enabled by `FALAFL` substantially helps identify CpG sites that are universally lineage-informative. For this purpose, we first perform the random selection of CpG sites from the left colon CRC subcohort input matrix for `FALAFL` as described in Section 3.1. Specifically, we randomly choose 1,346,130 sites to match with the number of sites chosen by `FALAFL` for the left colon CRC subcohort, and we repeat the random sampling 5 times, as detailed in Section 2.5. As can be observed in Figure 5b and the right panel of Figure 5a, the Pearson correlation of the JS distance of randomly selected sites are consistently lower than that of `FALAFL`-selected sites for each patient pair. In the left panel, it is also observed that the median distance(*ℓ, V*) of the JS distance of randomly selected sites are consistently higher than that of `FALAFL`-selected sites.

**Figure 5:**
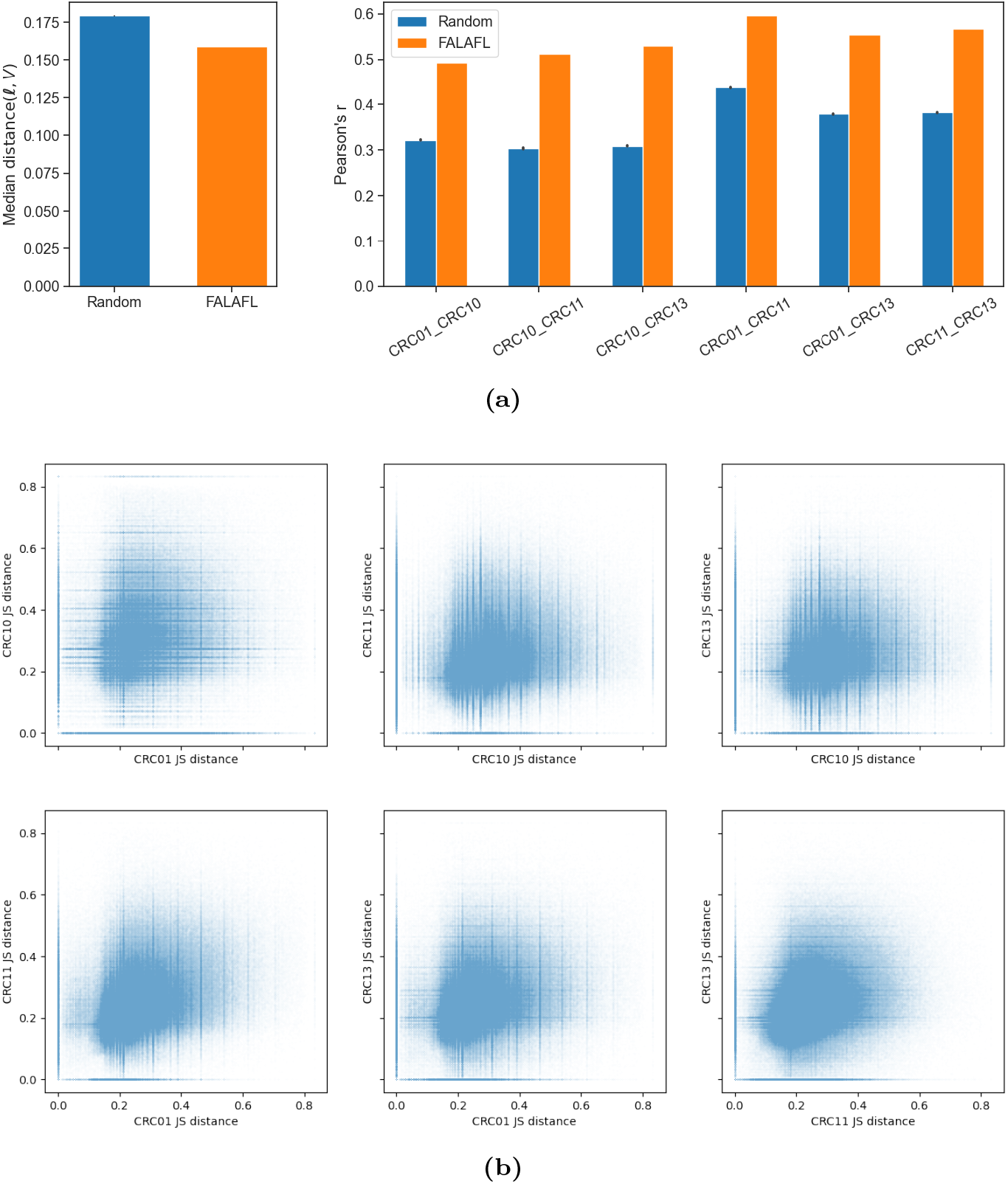
Benchmarking FALAFL against random CpG site selection. **(a)** Comparing random site selection and `FALAFL`. An equal number of sites in the `FALAFL` output are randomly selected from `FALAFL` input matrix of the left colon CRC subcohort. This random selection is repeated 5 times. The left panel shows the median distance(*ℓ, V*) (i.e. deviation from perfect correlation) among the randomly selected sites vs that of the `FALAFL`-selected sites. The right panel shows the Pearson correlation of randomly selected sites vs the `FALAFL`-selected sites for each patient pair. Both panels demonstrate that `FALAFL` selected sites better support the universality of lineage informativeness than the randomly selected sites. **(b)** Correlation of lineage-informativeness of randomly chosen CpG sites across each pair of left colon cancer patients. See Figure 3a for comparison.

We also compare the universality of lineage-informativenes for CpG sites selected using the pairwise greedy approach against `FALAFL`-selected CpG sites. Interestingly, `FALAFL`-selected CpG sites exhibit consistently higher correlation of lineage-informativeness (evaluated by Pearson correlation) across all patient pairs in the entire CRC cohort. As demonstrated in Figure 4b, among the 36 pairs of 9 patients, all but 2 patient pairs have increased Pearson correlations of lineage-informativeness for `FALAFL`-selected CpG sites for the entire CRC cohort of 9 patients than sites selected using the greedy pairwise approach. The CpG sites selected by `FALAFL` for the left colon CRC cohort of 4 patients also demonstrate increased level of correlation of lineage-informativeness, though the improvement is trivial.

These benchmarking results demonstrate that the `FALAFL`-selected CpG sites consistently exhibit a higher level of universality for lineage-informativeness compared to sites chosen using the random approach or the greedy pairwise approach. Our results signify the role of fair molecular feature selection enabled by `FALAFL` in pinpointing CpG sites with universally consistent behavior.

### 3.5 FALAFL-identified universally lineage-informative and uninformative CpG sites reveals non-stochasticity of CpG methylation alterations in colorectal cancer progression

To understand the molecular and functional implications of universally lineage-informative and lineage-uninformative CpG sites, we stratify CpG sites based on their universal lineage-informativeness in each patient pair, as defined in Section 2.4. We identify universally lineage-informative and uninformative sites for both the left colon CRC subcohort of 4 patients and the entire CRC cohort of 9 patients. While the remaining CpG sites lacking universality in their behaviors across patients may also harbor interesting biological insights, this paper focuses on characterizing CpG sites with universal lineage-informativeness and uninformativeness to shed light on the commonality of methylation changes in colorectal cancer progression.

In the left colon CRC subcohort of 4 patients, `FALAFL` selects 13,46,130 sites. As shown in Figure 6a, among all the selected sites, the mean value of mean JS distances across all patients is 0.2224, and the mean deviation from perfect correlation is 0.1671. Based on the definitions in Section 2.4, 321,006 sites are universally lineage-informative and 384,937 universally uninformative in the left colon CRC subcohort.

**Figure 6:**
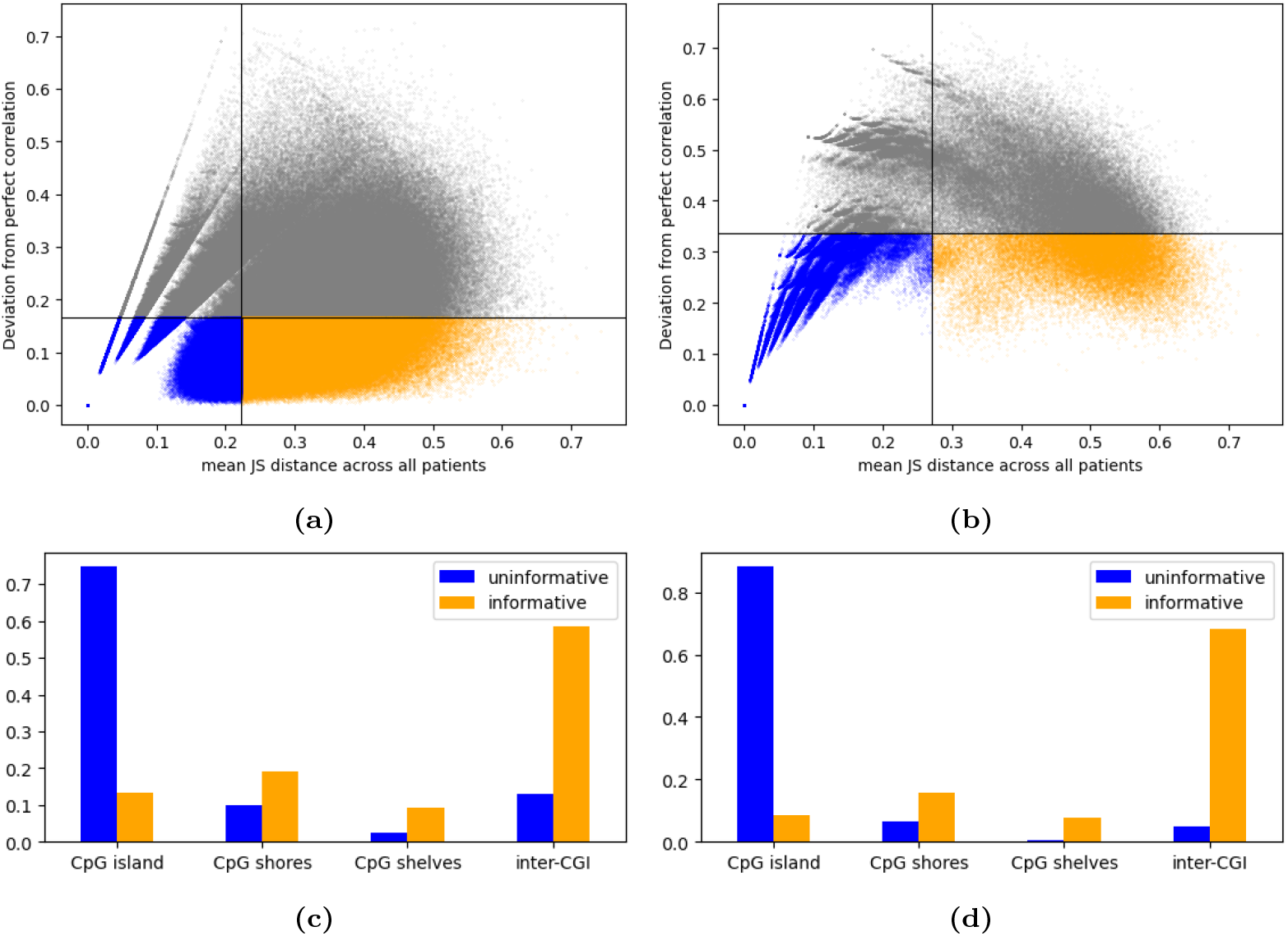
Characterization of FALAFL-identified universally lineage-informative and uninformative CpG sites in the left colon CRC subcohort and the entire CRC cohort. Universally lineage-informative sites are colored in orange, while universally lineage-uninformative sites are colored in blue. The remaining sites are colored in grey. **(a)** Scatter plot showing mean JS distance against deviation from perfection correlation of each `FALAFL`-selected CpG site for the left colon CRC subcohort of 4 patients. The vertical line marks the mean of mean JS distance across all patients, which is 0.2224. The horizontal line marks the mean deviation from perfect correlation, which is 0.1671.**(b)** Scatter plot showing mean JS distance against deviation from perfection correlation of each `FALAFL`-selected CpG site for the entire CRC cohort of 9 patients. The vertical line marks the mean of mean JS distance across all patients, which is 0.2717. The horizontal line marks the mean deviation from perfect correlation, which is 0.3355. **(c)** Bar plot showing the fractions of universally lineage-informative and uninformative CpG sites in left colon CRC subcohort located in CpG island, CpG shore, CpG shelf, and inter-CGI regions. **(d)** Bar plot showing the fractions of universally lineage-informative and uninformative CpG sites in left colon CRC subcohort located in CpG island, CpG shore, CpG shelf, and inter-CGI regions.

In the CRC patient cohort of 9 patients, `FALAFL` selects 195,809 sites to represent the entire cohort. As indicated in Figure 6b, among the 195,809 sites, the mean value of mean JS distances across all patients is 0.2717 and the mean deviation from perfect correlation is 0.3355. In the entire CRC patient cohort, 31,378 sites are deemed universally lineage-informative and 76,189 universally uninformative

Next, we categorize the universally lineage-informative and uninformative sites based on their genomic locations. Specifically, we map the sites to CpG islands (CGI, regions where CpG sites are over-represented, Cross and Bird 1995), CpG shores (2kb-long regions flanking both ends of CGIs, Irizarry et al. 2009), CpG shelves (2kb-long regions outside of CpG shores Irizarry et al. 2009), and inter-CGI regions. As demonstrated in Figure 6c and 6d, it is worth noting that in both the left colon CRC subcohort and the entire CRC cohort, universally lineage-informative sites are highly enriched in inter-CGI regions, whereas universally uninformative sites are predominantly localized in CpG islands. These results elucidate the non-stochastic nature of methylation changes in colorectal cancer progression and attest to the ability of `FALAFL`-selected molecular features to reveal interesting biological insights. It is possible that the universally lineage-informative sites identified by `FALAFL` can be further leveraged to gain insights into driver methylation events in cancer.

## 4 Conclusion

In this study, we present `FALAFL` (FAir muLti-sAmple Feature seLection), a novel fair molecular feature selection scheme aimed at ensuring algorithmic fairness in comparative patient profile analysis. Applied to the problem of fair selection of CpG sites within a cohort of colorectal cancer patients, our results demonstrate the ability of `FALAFL` in rapidly and robustly identifying a maximal set of CpG sites that are well-represented across the entire patient cohort. Importantly, `FALAFL` ensures fair representation of molecular features across all patients while uncovering cohort-wise universal behaviors of methylation changes. Furthermore, our investigation into the `FALAFL`-selected sites reveals interesting enrichment of universally lineage-informative CpG sites in inter CpG-island regions, which indicates a degree of non-stochasticity of methylation changes in colorectal cancer progression. These findings demonstrate the ability of `FALAFL` in identifying lineage-informative sites and providing valuable insights into the underlying biological mechanisms of cancer progression. We aim to apply `FALAFL` to larger and more diverse patient cohorts and further explore its effectiveness and generalizability in clinical settings.

## Footnotes

1 Note that read depth information is very different from methylation status information

2 While the Bian *et al*. cohort consist of 12 patients in total, we choose to exclude patients CRC03 and CRC06 for not having scBS-seq data and CRC09 for not having metastasis cells with scBS-seq data.

