## Supplementary Material for "Fair molecular feature selection unveils universally tumor lineage-informative methylation sites in colorectal cancer"

### S1 Code Availability

The code of `FALAFL` is available at Github (https://github.com/Storyboardslee/FALAFL).

### S2 Additional information of metastatic CRC patients from Bian *et al.* data set

| Patient | Primary site | Sampled lesions | Number of cells |
|---|---|---|---|
| CRC01 | Left Colon | NC, PT, LN, ML, MP | 160 |
| CRC02 | Right Colon | NC, PT, ML | 39 |
| CRC04 | Right Colon | NC, PT, LN | 65 |
| CRC10 | Left Colon | NC, PT, LN | 114 |
| CRC11 | Left Colon | NC, PT, LN | 228 |
| CRC12 | Rectum | NC, PT, LN | 22 |
| CRC13 | Left Colon | NC, PT, LN | 185 |
| CRC14 | Rectum | NC, PT, LN | 29 |
| CRC15 | Right Colon | NC, PT, LN, ML, MO | 50 |

**Table S1: Data summary of the Bian *et al.* metastatic CRC cohort [Bian et al., 2018].**

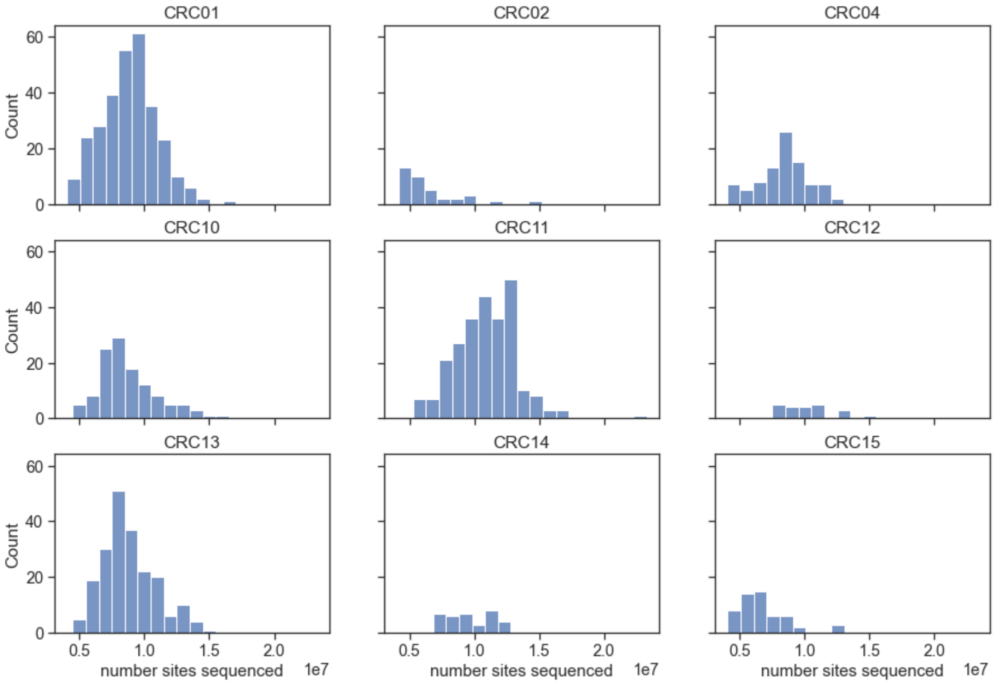


**Figure S1: Distribution of CpG site coverage of the Bian *et al.* metastatic CRC cohort.** The distribution plots shows the number of CpG sites sequenced in each cell on the x-axis, and number of cells on the y-axis.
